# Aerial survey perspectives on humpback whale resiliency in Maui Nui, Hawai’i, in the face of an unprecedented North Pacific marine warming event

**DOI:** 10.1101/2022.10.12.511738

**Authors:** Joseph R. Mobley, Mark H. Deakos, Adam A. Pack, Guilherme A. Bortolotto

**Author notes:** **Correspondence:** Joseph R Mobley, Jr., Dept. of Nursing, 2528 McCarthy Mall, University of Hawai‘i at Mānoa, Honolulu, Hawai‘i 96822. **Author contributions:** Conceptualization: J.M., A.P., M.D., Formal analysis: G.B., Funding acquisition: J.M., A.P., R.B. Investigation: J.M., A.P., M.D., Project administration: M.D., Writing—original draft: J.M. Writing—review and editing: A.P., M.D., G.B.

## Abstract

After decades of population growth, the central stock of the North Pacific population of humpback whales, known as the Hawai’i Distinct Population Segment (HDPS), was delisted from its endangered status in 2016. At that time, however, an unprecedented heating event, the “Pacific Marine Heatwave” (PMH) was already underway. The PMH coincided with reports of major declines of sightings of humpback whales, including calves of the year, on both the Hawaiian wintering grounds and the feeding grounds of Southeast Alaska. To examine the resiliency of the Hawai’i Distinct Population Segment, we conducted aerial surveys of the high-density Maui Nui region immediately following the PMH event in 2019 and 2020, using distance sampling methods identical to those used in an earlier series (1993 to 2003). Results showed whale densities at or above those seen earlier, with mean density for 2020 highest overall. Crude birth rates (% groups containing a calf) were similarly comparable to those recorded in the earlier series, with an increase from 2019 to 2020. Overall, results suggest the central North Pacific humpback whale population stock to be resilient in the face of this major climatic event.

## Introduction

After decades of international protection from commercial whaling since 1966 (Gambell, 1967), the population of North Pacific humpback whales has shown clear signs of recovery. Based on abundance estimates derived from over 18,000 identification images of individual humpback whales obtained from breeding and feeding regions throughout the North Pacific in 2004-06, and the “SPLASH” (Structure of Populations, Levels of Abundance and Status of Humpback Whales in the North Pacific) project, Barlow et al. (2011) concluded that, “the overall humpback whale population in the North Pacific has continued to increase and is now greater than some prior estimates of pre-whaling abundance” (p. 3). Other results from the same project showed Hawai’i to have the largest proportion of whales among the North Pacific breeding areas, representing 57% of that population, with an estimated abundance of approximately 10,000 individuals (Calambokidis et al., 2008). Abundance estimates across the three-year period of the SPLASH project showed an average annual rate of increase of 6%. This aligns with the earlier estimate of 7% increase per year for the main Hawaiian Islands obtained from fixed wing aerial surveys conducted across a ten-year period from 1993-03 (Mobley, 2004).

In September of 2016, considering the full suite of evidence for recovery, NOAA revised the status of the world’s populations of humpback whales and divided them into 14 Distinct Population Segments (DPS), removing nine from the endangered species list, listing four as endangered and one as threatened (Federal Register, 2016). The so-called “Hawai’i DPS” was one of those delisted from its initial 1970 endangered status under the U.S. Endangered Species Conservation Act. Ironically, during the two winter seasons just prior to this change in status (*i*.*e*., 2013/14 and 2014/15), the Northeast Pacific witnessed the Pacific Marine Heatwave (PMH), that was unprecedented in scope in recorded history (DiLorenzo & Mantua, 2016). The PMH lead to a cascade of environmental effects including increased sea surface temperatures, decreases in sea surface winds, less upwelling, a decline in nutrient rich water, and reduced phytoplankton and zooplankton, the foundation of the marine food chain (Gentemann, 2017).

Subsequent to the PMH, sharp declines in aggregations of humpback whales wintering in Hawaiian waters off west Maui, including calves-of-the-year, were reported for the years 2016-2018 based on visual transect surveys (Cartwright et al., 2019; Kügler et al., 2021), passive acoustic monitoring (PAM) in the ‘Au‘au Channel (Kügler et al., 2020, 2021), and from shore-based surveys off the Kona coast of Hawai‘i Island (Frankel et al., 2022). Both Cartwright et al. (2019) and Frankel et al. (2022) modeled the effects of various ocean climate indices including cycles of El Niño and Pacific Decadal Oscillation (PDO) on whale abundance and calf production in their respective study areas. Cartwright et al. (2019) found that mean encounter rates from vessel surveys decreased dramatically: by 76.5% for groups containing a calf and by 39% for non-calf groups; and also reported that a two-year lag in PDO explained approximately 67% of the variability in encounter rate data of calf groups. Frankel et al. (2022) reported that, during shore-based scans, whale numbers (*i*.*e*., average of whales per scan) increased over seasons, from 2001-2015, but after 2015 whale numbers decreased to their lowest level since 2000 and remained at low levels through 2019. Similarly, Frankel et al. (2022) reported that the annual mean crude birth rate dropped from 6.5% for 2001-2015 to 2.1% for 2016-2019). Statistical modeling revealed that both number of whales and crude birth rate declined during those breeding seasons, in which prior seasons on high latitude North Pacific feeding grounds revealed warmer waters based on indices of climate change. Likewise, greater numbers of whales observed on the Hawaiian breeding grounds followed negative PDO values (*i*.*e*., cooler, productive waters at high latitude feeding grounds) lagging by 1.5 years (*i*.*e*., the next breeding season). Modeling of crude birth rate in relation to climate factors was less clear. The best model indicated the crude birth rate increased in association with a 2.5-year lag from climate indices favoring productivity on summer feeding grounds, while the second and third best models indicated a 0.5-year lag.

Similar declines in humpback whale numbers were reported in feeding grounds of the Hawai’i DPS in southeastern Alaska including Glacier Bay, but starting as early as 2013 (Gabriele et al., 2022). In these areas, both non-calf and calf abundance decreased between 2013 and 2018, and then appeared to rebound in 2019-2020. The time lag in onset of population impact between the Alaskan and Hawaiian regions is consistent with the view that the PMH (augmented by a strong El Niño and positive PDO) likely negatively impacted humpback whale prey availability on the feeding grounds first, which may have in turn affected migratory arrivals and/or reproductive readiness of humpbacks on the Hawaiian wintering grounds (Frankel et al., 2022; Gabriele et al., 2022). Collectively these studies suggest that, despite the recovery of the Hawai’i DPS from commercial whaling (Barlow et al. 2011; Calambokidis et al. 2008), it remains fragile and susceptible to natural and anthropogenic threats, requiring continued monitoring, research and protection.

Despite these concerns, several authors have noted the apparent resilience of migratory baleen whales in the face of climate change. For example, Moore and Reeves (2018) explained that the migratory patterns of some Arctic baleen whales gave them an advantage over species dependent on ice shelves (*e*.*g*., polar bears), and that the extended periods of open water (2-4 weeks) due to global warming likely benefited them as well. In that work, the authors proposed that greater resilience arose from behavioral flexibility and resistance to disease and stress, assuming resilience as the ability to adapt to change. Derville et al. (2019) similarly remarked upon the relative resilience of humpback whales in particular owing to their “apparent plasticity of habitat use patterns” (p. 1466).

Here, we investigate the long-term resiliency of humpback whales taking up temporary residency on the Hawaiian breeding grounds in the broad areas of the “Maui Nui” region (encompassing the islands of Maui, Moloka‘i, Lāna‘i, and Kaho’olawe) by comparing the results of aerial surveys conducted from 1993-2003 (pre-MPH) with those from 2019-2020 (post-MPH). Based on earlier surveys of waters adjoining the main Hawaiian Islands, 63% of all whale sightings and 70% of all calves were seen in this region (Mobley et al., 1999).

## Methods

### Survey Periods

Aerial surveys consisted of an earlier series (1993-2003) flown during the years 1993, 1995, 1998, 2000, and 2003, as well as a later series (2019-2020) flown during the years 2019 and 2020. Survey design and data collection closely followed that of Distance Sampling methods (Buckland et al., 2001).

### Survey areas and transect placement

The 1993-2003 survey series consisted of aerial surveys around all of the main Hawaiian Islands including the Maui Nui region, incorporating the channels around the islands of Maui, Moloka‘i, Lāna‘i and Kaho‘olawe as well as Penguin Bank, which extends 25 nm southwest from the western end of Molokai. During the humpback whale breeding season, these areas host the greatest concentrations of humpback whales including calves of the year (Herman et al. 1980; Mobley et al. 1999). The 2019-2020 survey series was restricted to the Maui Nui region. All surveys followed north-south systematic lines placed 26 km apart with random legs connecting the endpoints (Figure 1). The north-south lines extended 13 km past the 1,828 m isobath (1000 fathoms) which occurred at an average distance of approximately 46 km offshore. The exact placement of lines varied on each survey by using random longitudinal start-points for the first survey of a given series.

**Figure 1.**
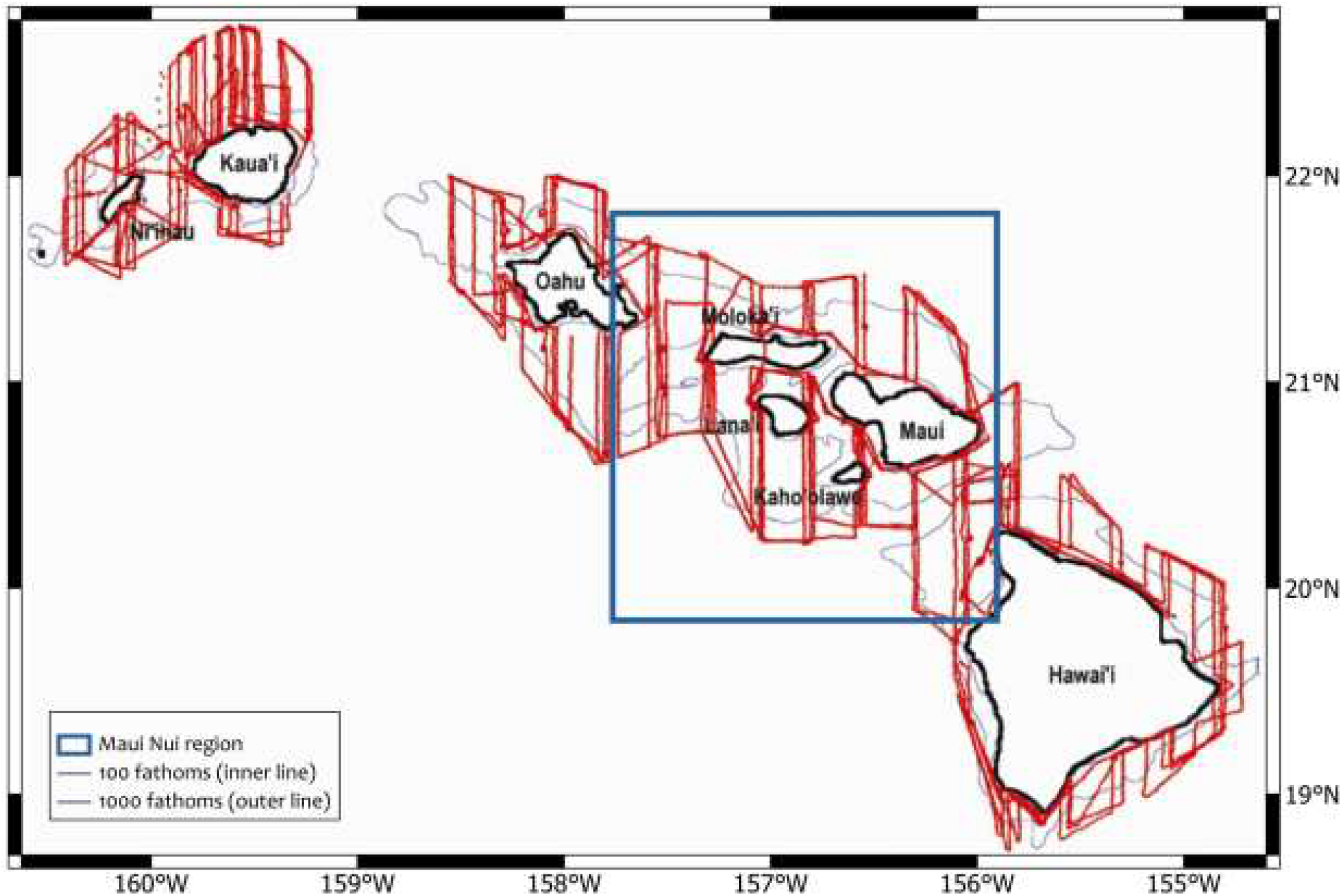
1993-2003 Survey Series. Surveys during each of the five years (2003 flight paths shown in red) covered waters adjoining all eight main islands. North-south systematic lines were placed randomly so the exact configuration of track lines varied across years with each extending 7 km past the 1,825 m (1,000 fathoms) isobath (light gray outer contour lines) (Figure adapted from Mobley, 2004).

### Data collection procedures

The survey aircraft was a fixed wing twin-engine Partenavia (P68) Observer flying at a mean speed of 100 knots and a mean altitude of 244 m. All crew wore internally connected headsets equipped with earphones and microphones to ensure clear communication. Two experienced observers on each side of the aircraft, equipped with Suunto brand hand-held clinometers, made sightings of humpback whales as well as any other marine mammal species. The windows through which each observer scanned allowed for 180-degree views in the horizontal plane and approximately 70-degree downward views, with a blind spot directly under the plane. For each sighting, observers reported the species, number of individuals present, presence or absence of a calf, and angle to the sighting (measured with the clinometer) when the sighting was abeam of the aircraft. A data recorder, seated next to the pilot, entered sighting information into an iPad with a customized Filemaker Pro database. A timestamp, GPS location, and altitude of the aircraft were collected from a Bluetooth-connected GPS (Bad Elf Pro) and automatically appended to each sighting entry. Additionally, GPS locations and altitude were automatically recorded every 30-seconds. Environmental data (Beaufort sea state and percent glare) were recorded at the start of each transect leg and whenever conditions changed. Perpendicular distances to each sighting were calculated using sighting angles combined with altitude data.

### Flight schedules

For the 1993-2003 series, a complete survey involved coverage of all eight major islands of the main Hawaiian chain (Figure 1), which required four days on average. Within each year, a total of four complete surveys of all the island regions were carried out spanning the period from mid-February through end of March, when past surveys have shown humpback whales to be most prevalent in the area (Herman and Antinoja, 1977; Baker and Herman, 1981; Mobley, Bauer and Herman, 1999).

### 2019 and 2020 Surveys

Surveys flown during the 2019-2020 series followed the same transect placement rules and data collection procedures as the earlier 1993-03 series, with two primary departures: a) the survey area was limited to the Maui Nui region (Maui, Moloka’i, Lāna’i, Kaho’olawe and Penguin Bank) (Figure 2) rather than all main islands; and b) three flights were flown in each year rather than four. As a result, comparisons of results from the earlier to the later series were limited to the Maui Nui region only.

**Figure 2.**
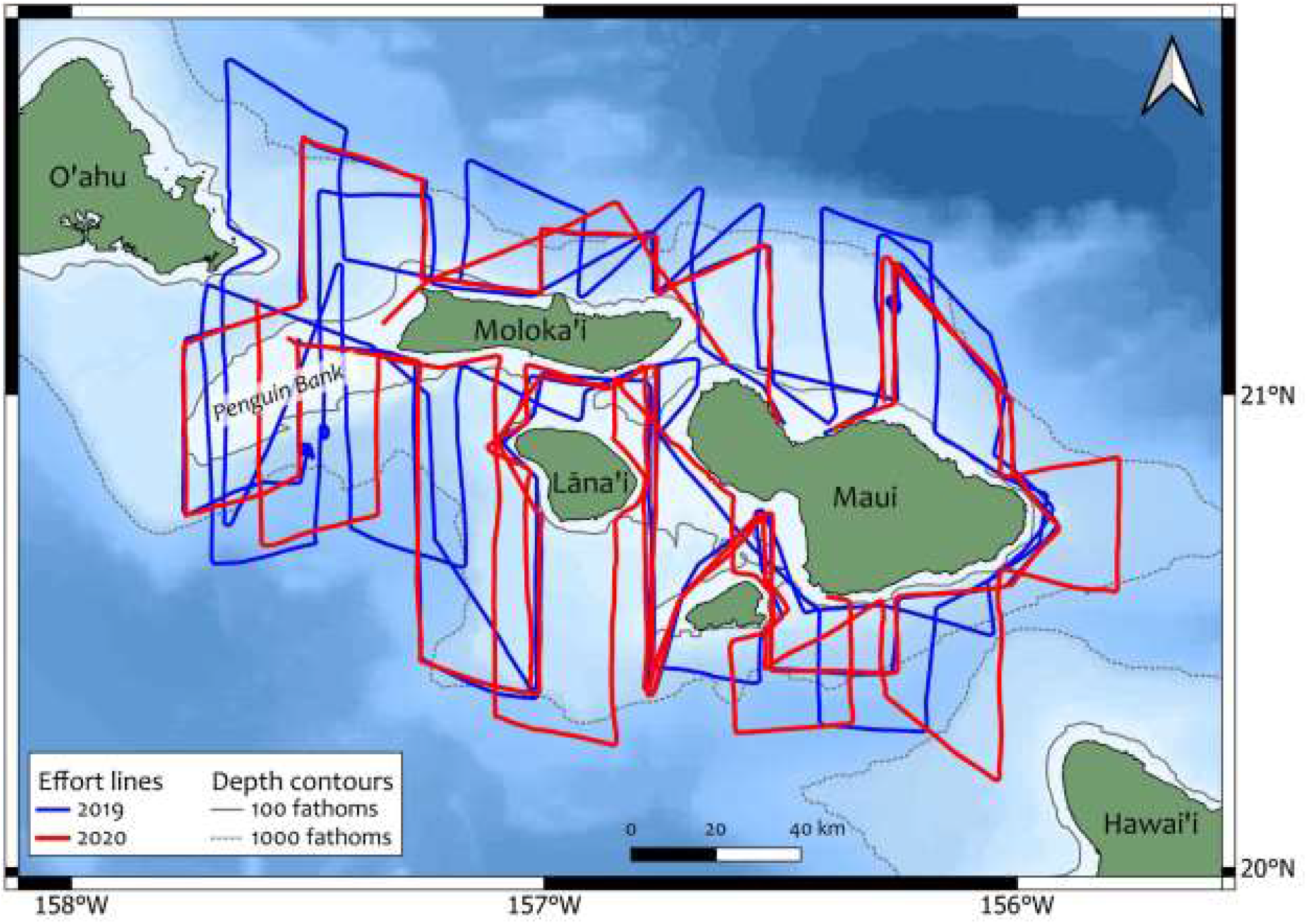
2019-2020 Survey Series. Later surveys covered waters within the Maui Nui region only (area within the square in Figure 1). Colored tracks represent survey lines flown on-effort in each year. The map background, representing bathymetric information, was created from data downloaded from http://www.soest.hawaii.edu/HMRG/multibeam/index.php. The 183 m (100 fathoms) isobath is demarcated by the gray solid line and the 1,825 m (1,000 fathoms) isobath is demarcated by the gray dashed line.

### Density estimates

Analyses were performed using software Distance version 7.3 and followed guidelines in Thomas et al. (2010). An exploratory analysis was conducted to decide appropriate truncation distances and to investigate the need for grouping data in distance intervals; that was followed by detection probability modeling and model selection. The last steps included final analysis and inferences. Since the Partenavia P-68 survey aircraft was outfitted with flat rather than bubble windows, viewing sightings directly underneath the aircraft was not possible. Thus, distances less than 200 m were truncated to account for the resulting blind spot. To model the detection function, a maximum truncation distance was assessed by visually inspecting perpendicular distance histograms. Sightings far from the survey lines, and consequently less frequent in the dataset, were excluded. Model selection was conducted in a stepwise approach, using conventional (CDS; Buckland et al., 2001) and multiple covariate distance sampling (MCDS; Marques et al., 2003), starting with simple models and including one adjustment term or covariate at a time. Half-normal and hazard rate models were considered as the key function.

CDS models included the following adjustment terms: cosine, hermite polynomial and simple polynomial, and MCDS models considered covariates of sea state category (3 or lower or 4 or higher, in the Beaufort scale) and group size.

Model selection was performed using AIC (Akaike’s Information Criteria), taking note of candidate models presenting some statistical support, indicated by an AIC within two units to that of the model presenting the smallest value. Empirical variances, coefficients of variation (CVs), 95% confidence intervals (Satterthwaite’s approximation method; Buckland et al, 2001) were also estimated in software Distance. The selected model for detection function was used to estimate whale densities for each year (*i*.*e*., post-stratified by year). A global mean group size, considering the entire dataset after truncation, was used in the density estimation, as group size bias (Buckland et al., 2001) was verified not to be an important issue here. Since the present analysis ignored the “g(0) < 1 issue” (Buckland et al. 2001), which happens when the assumption of “animals over the survey line have a detection probability equal to 1” is not met, the estimates produced here are relative densities and should not be used to infer abundance. For a discussion of the requirements for estimating absolute abundance using distance sampling data from cetacean surveys, the reader is referred to Buckland et al. (2001).

## Results

### Effort and Sightings

Across a 27-year span (from 1993-2020), surveys took place during seven humpback whale breeding seasons, with a 15-year gap between the 1993-2003 and 2019-2020 series, when no surveys were flown (Table 1). Because the 1993-2003 series included all regions within the eight main islands of the Hawaiian Island chain, whereas the 2019-2020 surveys were specific to the Maui Nui region, for comparability, only the survey effort and humpback whale sightings within the Maui Nui region were included in the present analysis. All surveys combined for the Maui Nui region comprised a total of 30,265 km of linear effort, with 1,542 sightings (groups of whales) available for detection function analysis (Table 1) after distance truncation. That total was lower than the original number of sightings, since sightings beyond the truncation distance were ignored in the analysis. Effort covered shallow inland waters out to more pelagic areas beyond the 1,829 m (1000 fathom contour; Figures 1 and 2).

**Table 1.** Summary of effort and number of humpback whale sightings per year in the Maui Nui region, used in the density analysis.

| Year | Date range | Effort (km)* | No. lines | No. sightings |
| --- | --- | --- | --- | --- |
| 1993 | Feb. 21<sup>st</sup> - Mar. 26<sup>th</sup> | 5,551 | 159 | 238 |
| 1995 | Feb. 28<sup>th</sup> - Apr. 7<sup>th</sup> | 7,818 | 204 | 453 |
| 1998 | Feb. 21<sup>st</sup> - Apr. 17<sup>th</sup> | 4,061 | 119 | 180 |
| 2000 | Feb. 21<sup>st</sup> - Apr. 8<sup>th</sup> | 3,673 | 115 | 148 |
| 2003 | Feb. 21<sup>st</sup> - Apr. 5<sup>th</sup> | 4,175 | 124 | 203 |
| 2019 | Feb. 8<sup>th</sup> - Mar. 1<sup>st</sup> | 2,818 | 95 | 151 |
| 2020 | Feb. 13<sup>th</sup> - Mar. 2<sup>d</sup> | 2,170 | 79 | 169 |
| **Total** | - | **30,265** | **895** | **1,542** |

\*Note: The variability of linear effort across years represents two primary influences: a) effort for the 1993-2003 series involved four flights vs three flights for the 2019-20 effort; and b) depending on the random placement of the first north-south line, going 7 nmi past the 1,000 fathoms contour (per our trackline protocol) could produce broad variability in the length of those tracklines.

Humpback whale distribution patterns in 2019-20 (Figure 3) were similar to those described earlier (Mobley et al., 1999; Herman & Antinoja, 1977), *i*.*e*., showing a clear preference for shallow waters inside the 100-fathom isobath (64% of sightings in 2019, and 75% in 2020), mostly on the leeward sides of islands (86% in 2019, and 96% in 2020), with relatively few in the windward regions. The exception to the latter pattern is the high concentration of whales seen on the Penguin Bank, southwest of Molokai. Unlike the other high density areas, Penguin Bank is fully exposed to the trade winds with little protection from the surrounding islands. Sightings and effort data used in the analysis (*i*.*e*., after truncation) are summarized in Table 1.

**Figure 3.**
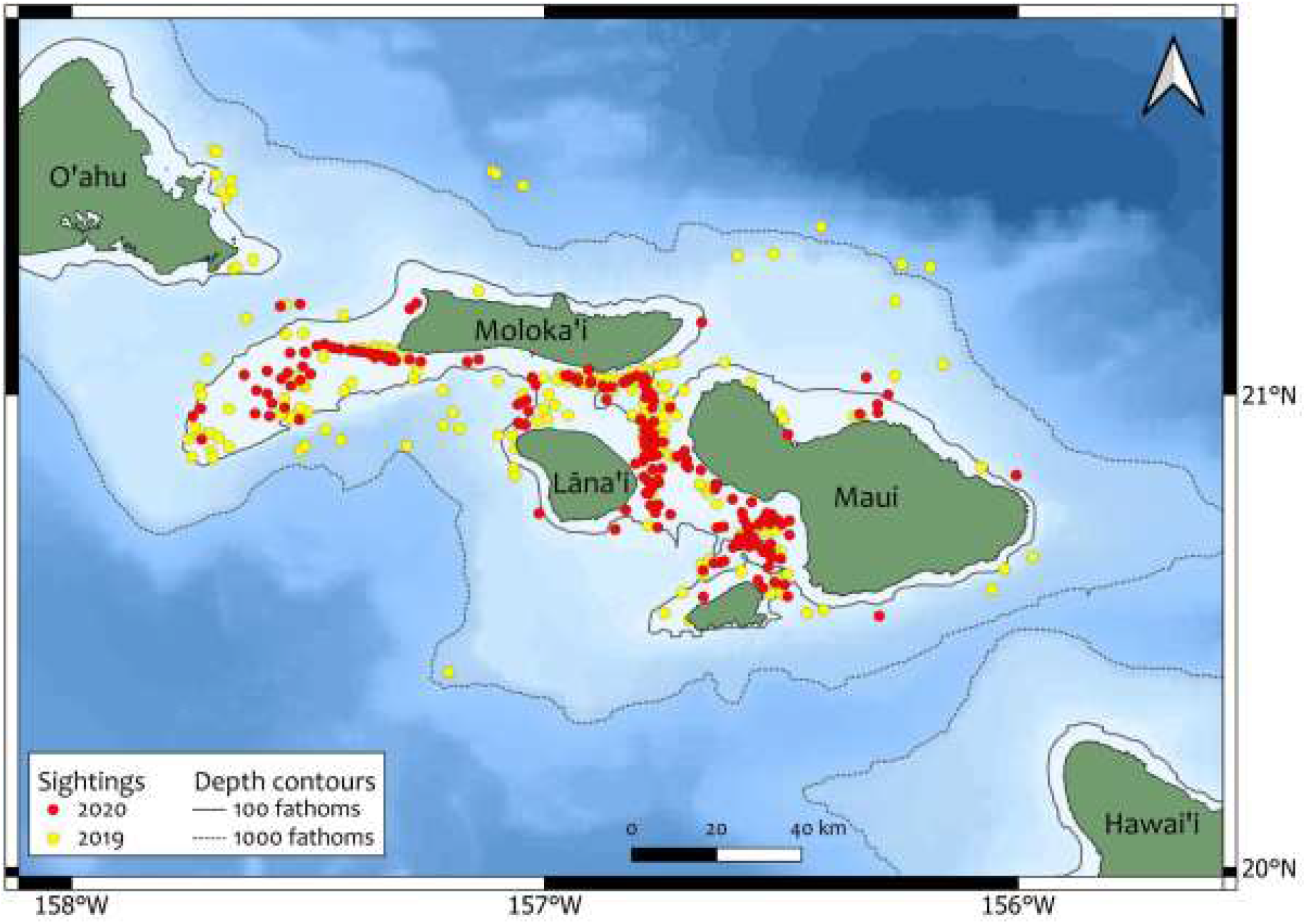
Distribution of humpback whale sightings during aerial surveys of the Maui Nui Region in 2019 and 2020. The map background, representing bathymetric information, was created from data downloaded from http://www.soest.hawaii.edu/HMRG/multibeam/index.php. The 183 m (100 fathoms) isobath is demarcated by the gray solid line and the 1,825 m (1,000 fathoms) isobath is demarcated by the gray dashed line.

### Density Estimation

A half-normal model with cosine adjustment (order 2) was selected for the detection function. Kolmogorov-Smirnov test results indicated a good fit (K-S = 0.0206, p = 0.532). A histogram of the estimated distances with fitted probability density function for the selected model is shown in Figure 4. The inclusion of a covariate sea state category in the model showed some support with a delta AIC equal to one (*Supporting information S1*) but ultimately the more parsimonious (*i*.*e*., with fewer parameters) model was chosen.

**Figure 4.**
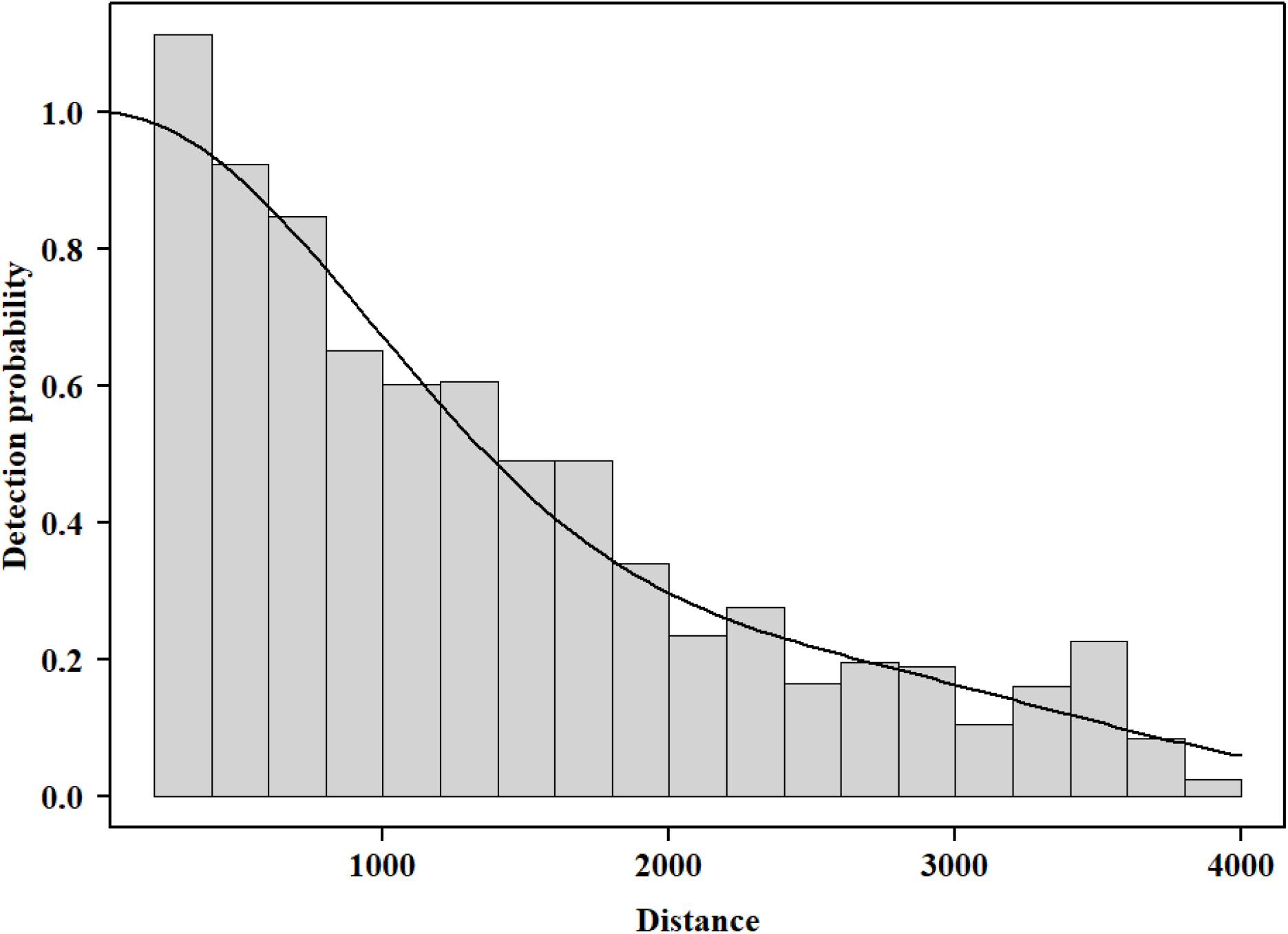
Perpendicular distances and fitted probability density function for all sightings (1993-2020).

Table 2 shows density results by year for each of the seven years surveyed. Density estimates for individual whales ranged from 0.022 (2000) to 0.043 (2020). There was also considerable uncertainty, with CVs ranging from 11.4% to 22.4%,with less precise density estimates particularly for 2019 and 2020.

**Table 2.** Density estimates of humpback whale groups and individuals in the year surveyed with coefficients of variation (CV) and confidence intervals (CI).

| Year | Category | Estimate | %CV | 95% CI |  |
| --- | --- | --- | --- | --- | --- |
| 1993 | groups | 0.015 | 14.7 | 0.011 | 0.019 |
|  | individuals | 0.024 | 14.8 | 0.018 | 0.032 |
| 1995 | groups | 0.020 | 11.3 | 0.016 | 0.025 |
|  | individuals | 0.032 | 11.4 | 0.026 | 0.040 |
| 1998 | groups | 0.015 | 20.1 | 0.010 | 0.022 |
|  | individuals | 0.025 | 20.2 | 0.017 | 0.036 |
| 2000 | groups | 0.014 | 22.8 | 0.009 | 0.021 |
|  | individuals | 0.022 | 22.9 | 0.014 | 0.035 |
| 2003 | groups | 0.017 | 21.8 | 0.011 | 0.025 |
|  | individuals | 0.027 | 21.8 | 0.018 | 0.041 |
| 2019 | groups | 0.018 | 17.7 | 0.013 | 0.026 |
|  | individuals | 0.030 | 17.8 | 0.021 | 0.042 |
| 2020 | groups | 0.027 | 22.4 | 0.017 | 0.041 |
|  | individuals | 0.043 | 22.4 | 0.028 | 0.067 |

Figure 5 shows the same data from Table 2 in graphic form, with a regression line fitted to all seven density points. However, the suggestion of an increasing trend is largely influenced by the density estimate for 2020, and the resulting linear trend was not statistically significant (p = 0.09). An additional investigation of the uncertainty for the positive trend is presented in *Supporting Information S2*.

**Figure 5.**
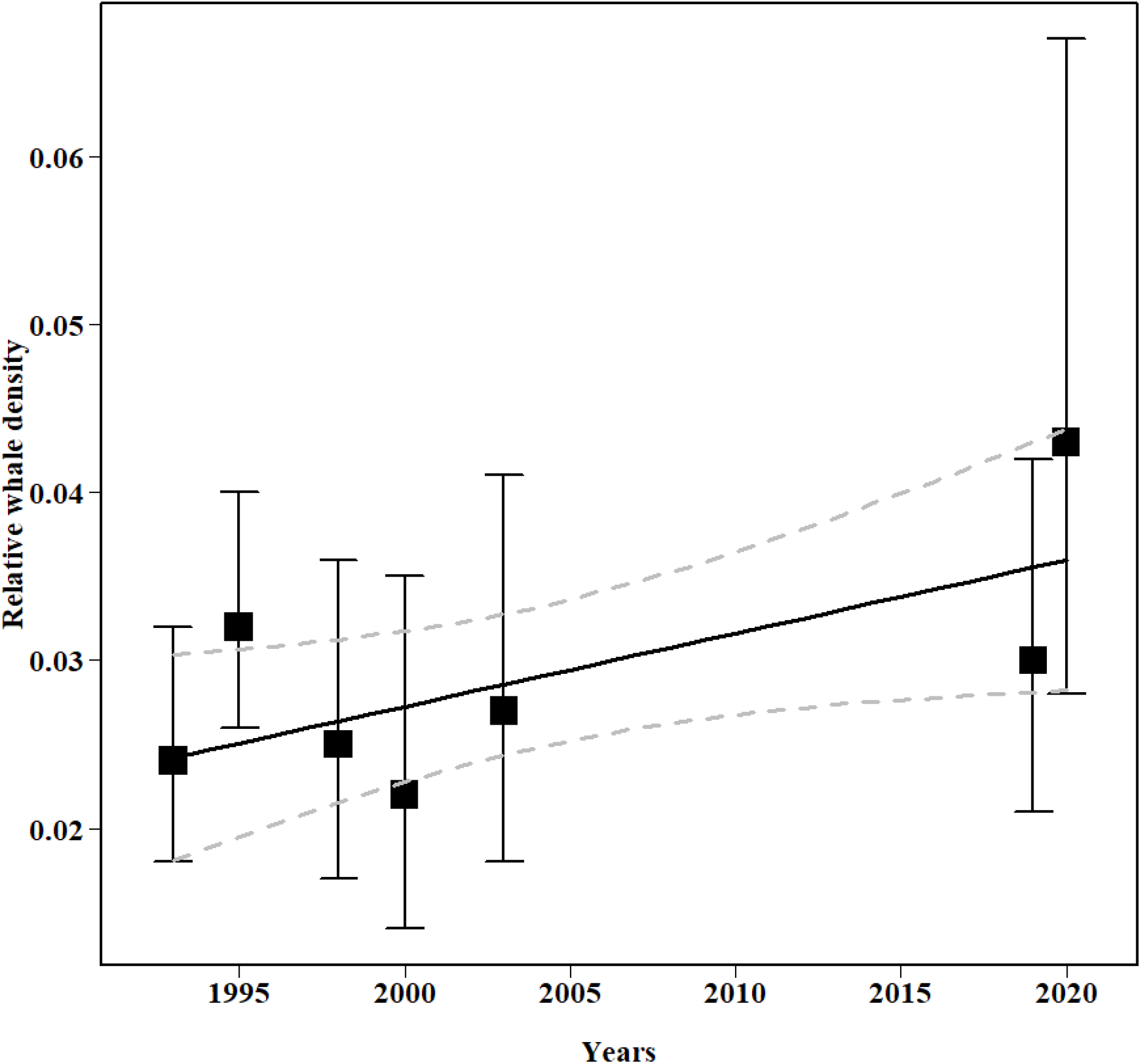
Humpback whale densities by year with a regression line fitted to the point estimates for individuals (Table 2). Though suggestive of a gradual increase, that trend has weak statistical support (p = 0.09) (see *Supporting information S2* for details on the uncertainty in the estimated trend). Dashed lines represent 95% percentile confidence intervals for the fitted regression).

### Crude Birth Rate

Other than changes in overall whale density, another indicator of overall population status is the proportion of whale groups that contained calves, or crude birth rate (e.g., Craig & Herman, 2000). The crude birth rate observed in the Maui Nui region ranged from 5% in 1995 to 10.5% in 2020, for an overall mean of 8.5% (Figure 6). The rates in 2019 and 2020 are comparable to those seen during the earlier survey series, suggesting that the wintering population is currently holding steady.

**Figure 6.**
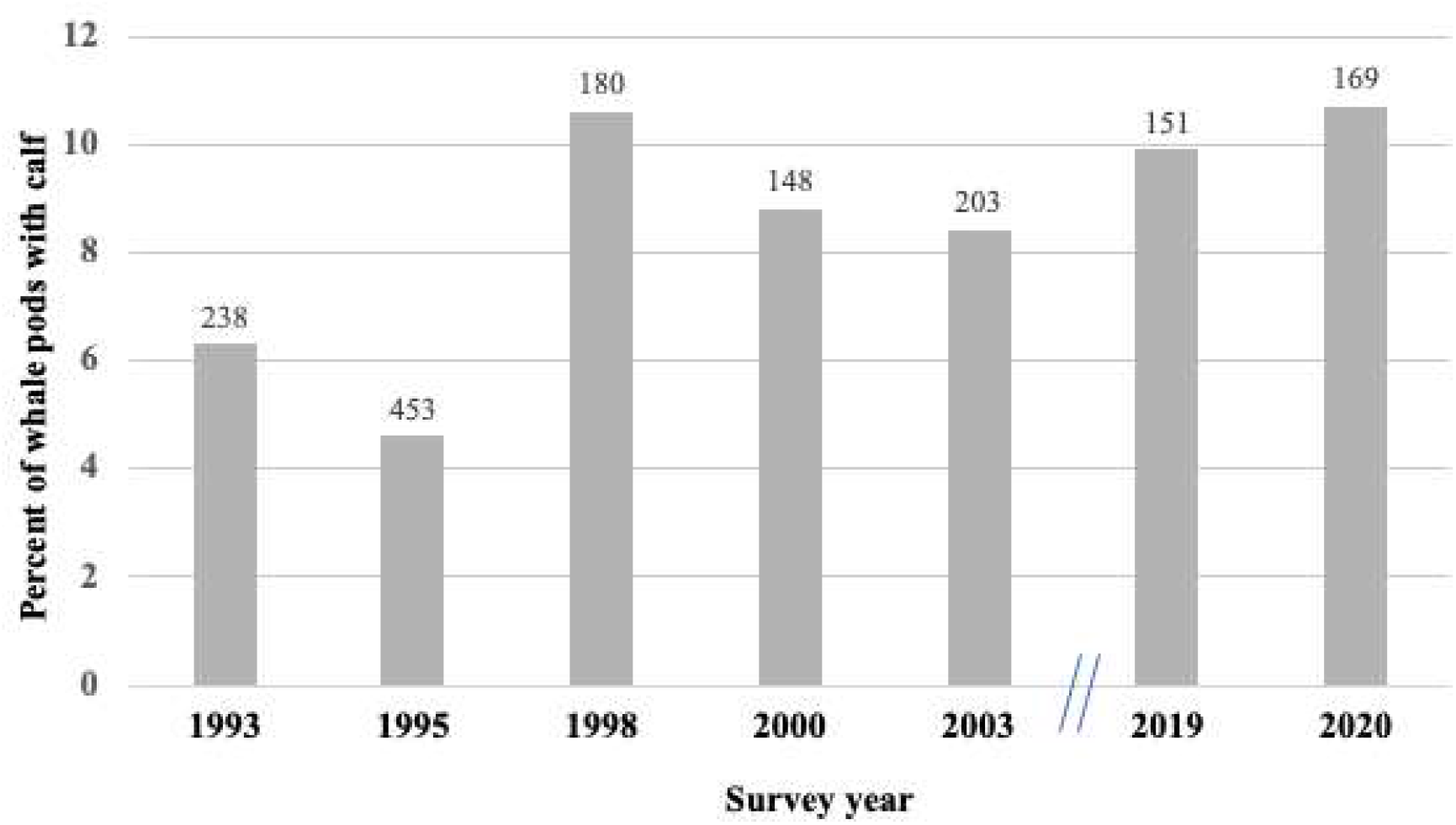
Percent of whale groups containing a calf by year with total sightings shown above each column.

## Discussion

Our findings provide a temporally and spatially broad perspective on the resiliency of the Hawai’i DPS of humpback whales in the face of an unprecedented North Pacific marine heatwave in the wintering area of historically the highest whale concentration. Temporally, our comparative analyses included a 15-year gap between early and later aerial survey series which is greater than previous reports examining the effect of the marine heatwave on humpbacks in Hawai’i waters using boat-based, shore-based and PAM survey techniques. We also covered a more expansive area in the four-island region than these previous studies.

### Density Trends

Our findings using aerial survey distance sampling techniques indicated that humpback whale densities in 2019-2020 were comparable with estimates from 1993-2003. Thus, despite an intervening marine heatwave from 2014-2017 coupled with overlapping El Niño and positive PDO marine warming events, which negatively impacted some North Pacific humpback whales’ food resources and resulted in observed reductions in whale and calf sightings in the Hawai’i DPS during these years by Cartwright et al. (2019), Frankel et al., (2022) and Kügler et al. (2020), the abundance and calf production of humpback whales in the Maui Nui region appears resilient. This finding is consistent with Kügler et al. (2020) who, based on PAM recordings at six sites in the Maui Nui region during the breeding seasons of 2014-2015, 2017-2018 and 2018-2019, found significant decreases in the acoustic energy which is dominated by humpback whale song (and reflective not only of male singers but of overall whale abundance; see Kügler et al. 2021), at all sites from 2014-15 and 2017-18 followed by increases at some of the sites during 2018-19. In the same region, Cartwright et al. (2019) conducting boat surveys from 2008-2018, found increases in encounter rates of humpback whales late in the breeding season (which encompassed the greatest span of annual surveys) from 0.42 whales/km in 2008 to 1.12 whale/km in 2013 with decreases thereafter to 0.14 whales/km in 2018. The declining trend in encounter rates from 2013 to 2018 was consistent when Cartwright et al. (2019) considered the full breeding season (early, mid and late season) surveys, either across individual years (i.e., 2013>2014>2017>2018) or the pairing of years 2013-2014 versus 2017-2018. Although Cartwright et al. (2019) did not report on boat surveys in 2019, Zang and Lammers (2021) conducted boat-based surveys using a more expanded footprint than Cartwright et al. (2019) and showed an increase in estimated peak abundance and density from the 2019 breeding season (winter 2018-spring 2019) to the 2020 breeding season (winter 2019-spring 2020), although with some fluctuation in the 2021 breeding season.

Using shore-based scans spread over the period of each day in the South of the Maui Nui region, off north Kona coast of Hawai’i Island, Frankel et al. (2022) showed a generally steady increase in the mean number of whales per scan from 2001-2015 (overall annual mean number of whales per scan = 19.2), and then a significant drop in 2016, with an overall annual mean number of whales per scan of 13.1 from 2016-2019. Although the mean number of whales per scan had a general positive slope from 2016-2019, showing perhaps the beginnings of recovery, the mean number of whales per scan in 2019 remained relatively low compared to all but one year, from 2009 through 2015.

The putative effects of the PMH event were not limited to the Hawaiian wintering grounds. Gabrielle et al. (2022, see Figure 3 in that article) described a similar pattern of decline for Icy Strait and Glacier Bay on the Southeast Alaskan feeding grounds, coinciding with the PMH event (2014-2016). Abundance levels reached their lowest point in 2018 followed by marked increases during both 2019 and 2020.

The PMH event caused die-offs and reproductive failures of several marine species including murres (*Uria aalge*) (Piatt et al., 2020). Piatt et al. described how the PMH caused metabolic increases at multiple trophic levels which resulted in an “ectothermic vise” mechanism. Presumably these disruptions in the trophic chain affected prey availability at higher levels including the prey species for humpback whales. This in turn potentially affected their readiness for migration and/or reproduction in general.

### Crude Birth Rate

The crude birth rate (CBR) estimates presented here for the years immediately following the PMH event were 9.2% and 10.5% for 2019 and 2020 respectively. These estimates correspond to those from our aerial surveys prior to the PMH event (Figure 6). Nonetheless, either through inference based on encounter rates with calf groups or directly observed, the CBR was particularly impacted during and immediately following the years corresponding to the PMH event (2014-2017).Also, boat-based surveys off west Maui found for groups containing a calf a late season increase in encounter rates from 2008-2013 with a declining trend thereafter (Cartwright et al., 2019)). Comparing the breeding seasons of 2013-2014 to 2017-2018, encounter rates for groups containing a calf dropped 76.5%. Year-by-year, calf-groups encounter rates showed a precipitous decline from 2013 to 2014, 2017 and 2018. For the North Kona coast of Hawai‘i Island, Frankel et al. (2022)’s shore-based surveys showed that for 2001-2015, the mean CBR was 6.5% (range = 4.7-9.6), while in 2016-2019, it dropped to 2.1% (range = 1.1-2.9).

The observation that CBR estimates for the Kona coast remained low (2.1%) up to 2019, when those for the Maui Nui region were above 9% may represent sampling error and the use of different techniques (shore-based surveys versus aerial surveys), but it is consistent with the observation that Maui Nui and Hawai‘i Island represent habitats with different utilization patterns by mature females. For example, Craig and Herman (2000) studying humpback identification photographs for 1977-1994, noted that CBRs were higher for Maui than Hawai‘i Island, and that individual females seen in both regions in different years were more often accompanied by a calf when in Maui Nui than Hawai‘i Island. Thus, Maui Nui appears to be utilized by many females as a preferred area of calving compared to Hawai‘i Island. However, if there were significant die offs of humpback whales during the PMH that traditionally migrate to the Maui Nui Region, it is also possible that the continued low CBR for Hawai’i Island reflects more gravid females seizing an opportunity to take up residence in the Maui Nui region, instead of Hawai’i Island.

### Impacts of ocean climate changes on humpback whale abundance and calving in Hawaiian waters

Both Cartwright et al. (2019) and Frankel et al. (2022) modeled the effects of various ocean climate indices including cycles of El Niño and Pacific Decadal Oscillation (PDO) on measures related to whale abundance and calf production in their respective study areas. As noted earlier, Cartwright et al. (2019) found that mean encounter rates from vessel surveys decreased dramatically from 2013-2014 compared to 2017-2018, by 76.5% for groups containing calves, and by 39% for non-calf groups. They reported that a two-year lag in PDO explained approximately 67% of the variability in encounter rate data of calf groups. Frankel et al. (2022) reported that during shore-based scans, whale numbers (*i*.*e*., average of whales per scan) increased over seasons, from 2001-2015, but after 2015 whale numbers decreased to their lowest level since 2000 and remained at low levels through 2019. Likewise, between 2015 and 2016, Frankel et al. (2022) reported that the annual mean crude birth rate dropped from 6.5% (2001-2015) to 2.1% (2016-2019). Statistical modeling revealed that both numbers of whales and crude birth rate declined during those breeding seasons in which prior seasons on high latitude North Pacific feeding grounds revealed warmer waters based on indices of climate change. Similarly, greater numbers of whales on the Hawai’i breeding grounds were predicted by negative PDO values (*i*.*e*., cooler, productive waters at high latitude feeding grounds) lagging by 1.5 years (the next breeding season). Modeling of crude birth rate in relation to climate factors was less clear. The best model indicated the crude birth rate increased in association with a 2.5-year lag from climate indices favoring productivity on summer feeding grounds, while the second and third models indicated a 0.5-year lag. Clearly, the collection and analyses of more long-term time series of climate variability indices in conjunction with humpback whale abundance estimates, CBR and habitat areas in the Hawaiian Islands is necessary to refine these estimates of lag.

### Summary and Future Research Directions

When all available data summarized here are combined, the reported decline in numbers of both adult humpbacks and calves largely overlapping with the PMH event (2014-2016) provides compelling evidence of its impact on this recovering DPS, both for their wintering grounds in Hawai’i and their feeding grounds in Southeastern Alaska. However, based on the current results showing increases of both density and CBR estimates for 2019 and 2020, there appeared to be a rather quick recovery from the effects of the PMH. These increases are corroborated by the post-event data of Kügler et al. (2021) for 2019 based on SPLs of chorusing whales off west Maui (see Figure 6 in that article), as well as the 2019-2020 abundance estimates of Gabrielle et al. (2022) for southeastern Alaska. This provides evidence of resilience for this distinct population segment (DPS) in the face of climate change (Moore & Reeves, 2018; Derville et al., 2019).

A number of questions remain unaddressed, however. First, because formal surveys during and following the PMH were limited to the Maui Nui Region and Hawai’i Islands, it is unclear whether the PMH uniformly affected the humpbacks across the other main Hawaiian Islands and extended to the Northwestern Hawaiian Island archipelago. Conducting all-island aerial surveys as was done in the early series reported here and relating findings to various climate indices can provide important insight, especially given that habitat use in some island areas such as off Kauai has been relatively recent compared to other areas such as the Maui Nui Region (Mobley et al. 1999). Similar insight may be provided by continued PAM monitoring during the breeding season of waters off the Northwestern Hawaiian Islands which, because of accessibility and sea state during winter months, can be more challenging for boat-based or shore-based surveys. The humpback whale population of the latter region has only recently come under study (e.g., Lammers et al., 2011). A second question is about changes in residency during the PMH. Well prior to the PMH, Craig et al. (2001) found that the duration of residency was a function of sex, age-class and reproductive condition with mothers of newly born calves and mature males having the longest residency periods. It is unknown how residency characteristics were affected during the PMH, for example, whether mothers who must not only support themselves, but also their calves, based on metabolizing fat stores had to truncate their residency to a greater degree than mature males. Third, it is unknown whether whales that historically have long-term sighting histories on the Hawaiian breeding grounds (Herman et al. 2011) continued to migrate to Hawai’i even in the face of the PMH or whether these whales switched breeding grounds or simply overwintered on the feeding grounds. Finally, the limitations of the resilience of humpback whales to climatic change are yet to be elaborated as well. It is likely that climate events such as the PMH will continue to test the resilience of this sentinel species.

## Acknowledgments

We are grateful to our sponsors of the 2019-2020 aerial surveys including Dave Jung of Hawaii Ocean Project, Whale Trust, and The Dolphin Institute. We are also grateful to our excellent pilot, William “Billy” Zeffiro. Rick Camp provided useful insights into the uncertainty in the density estimates, as well as our additional observer Elia Herman. Aerial surveys were flown under NOAA permit number 21482 issued to Dan Engelhaupt.

