## Supporting information for "Aerial survey perspectives on humpback whale resiliency in Maui Nui, Hawai’i, in the face of an unprecedented North Pacific marine warming event"

**for**

**by**

**Mobley, Deakos, Pack and Bortolotto**

### S1 | Alternative detection function model including covariate "Beaufort category"

Model description: half-normal key with cosine adjustment (order 2) and covariate Beaufort category for the detection function

Delta AIC: 1.05

Kolmogorov-Smirnov test results: K-S = 0.0199, p = 0.573.

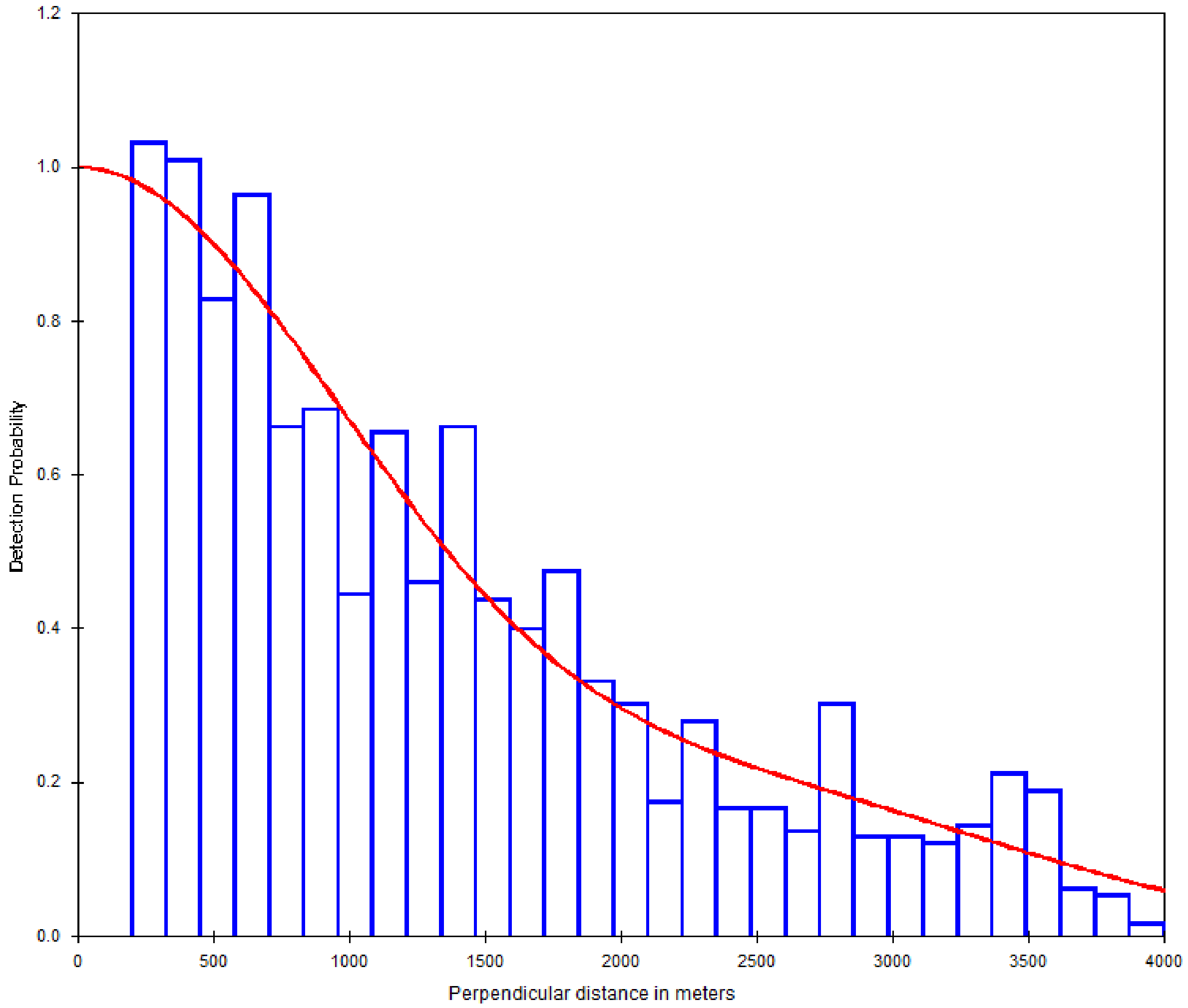


**Figure S1.1.** Detection function for a half-normal key with cosine adjustment (order 2) and covariate Beaufort category fitted to detection distances data

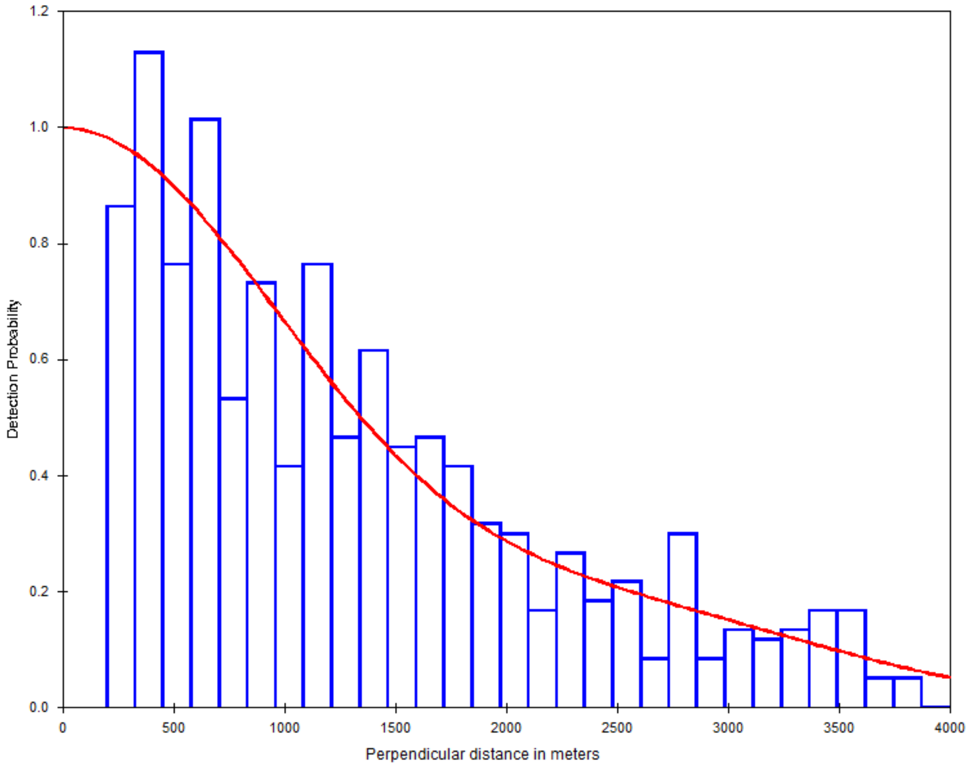


**Figure S1.2.** Detection function for a half-normal key with cosine adjustment (order 2) and covariate Beaufort category fitted to detection distances data, for detections made at sea state category "3 or lower" (in the Beaufort scale).

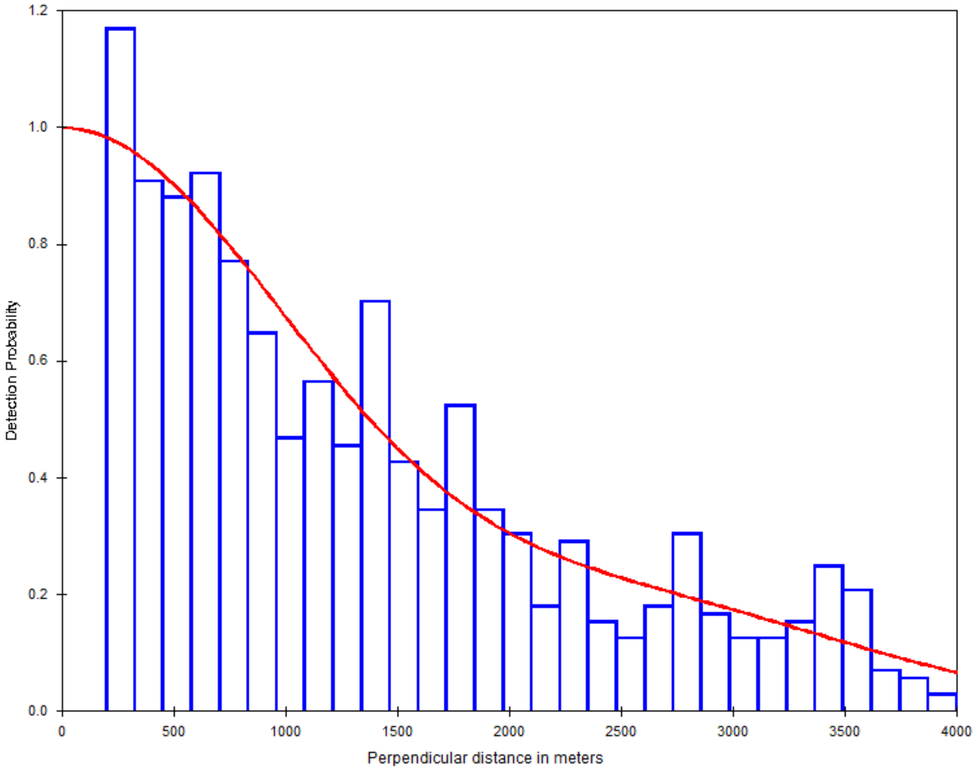


**Figure S1.3.** Detection function for a half-normal key with cosine adjustment (order 2) and covariate Beaufort category fitted to detection distances data, for detections made at sea state category "4 or higher" (in the Beaufort scale).

**Table S1.** Density estimates for a half-normal key with cosine adjustment (order 2) and covariate Beaufort category

| Year | Density | Estimate | %CV | 95% CI | |
|---|---|---|---|---|---|
| **1993** | **groups** | 0.015 | 14.4 | 0.011 | 0.019 |
| | **individuals** | 0.024 | 14.5 | 0.018 | 0.032 |
| **1995** | **groups** | 0.020 | 11.0 | 0.016 | 0.024 |
| | **individuals** | 0.032 | 11.1 | 0.026 | 0.040 |
| **1998** | **groups** | 0.015 | 19.9 | 0.010 | 0.022 |
| | **individuals** | 0.025 | 20.0 | 0.017 | 0.036 |
| **2000** | **groups** | 0.014 | 22.7 | 0.009 | 0.021 |
| | **individuals** | 0.022 | 22.7 | 0.014 | 0.035 |
| **2003** | **groups** | 0.017 | 21.6 | 0.011 | 0.025 |
| | **individuals** | 0.027 | 21.6 | 0.018 | 0.041 |
| **2019** | **groups** | 0.018 | 17.5 | 0.013 | 0.026 |
| | **individuals** | 0.030 | 17.5 | 0.021 | 0.042 |
| **2020** | **groups** | 0.027 | 22.2 | 0.017 | 0.041 |
| | **individuals** | 0.043 | 22.2 | 0.028 | 0.067 |

*Supporting Information S2*

**for**

**Aerial survey perspectives on humpback whale resiliency in Maui Nui, Hawai'i, in the face of an unprecedented North Pacific marine warming event**

**by**

**Mobley, Deakos, Pack and Bortolotto**

### S2| Uncertainty in density estimates trend

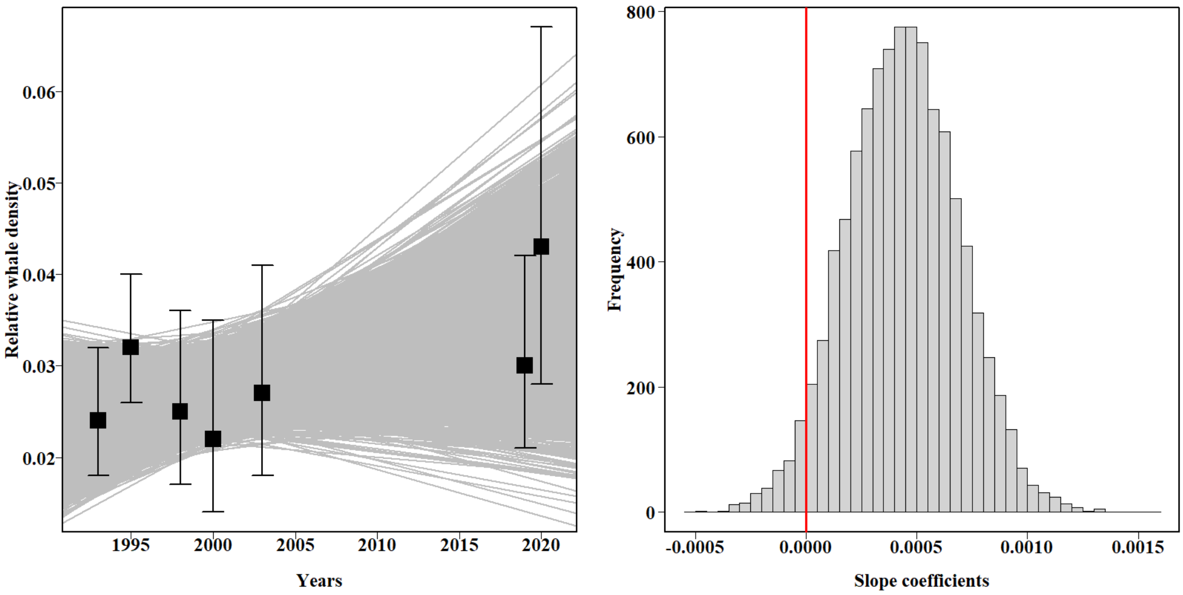


**Figure S2.** Right: the gray lines represent 10,000 linear models fitted to random draws from normal distributions centered at the point estimates of whale density (black squares), with the standard deviation derived from the corresponding coefficient of variation (Table 3 in the main text), for each year; Left: histogram of slope coefficients of 10,000 linear models with "zero" (vertical red line) highlighting the region in the graph that would represent stability; note that the large majority of the slopes indicate a positive trend (*i.e.*, > 0).
